# Nascent mutant Huntingtin exon 1 chains do not stall on ribosomes during translation but aggregates do recruit machinery involved in ribosome quality control

**DOI:** 10.1101/2020.05.13.094060

**Authors:** Angelique R. Ormsby, Dezerae Cox, James Daly, David Priest, Elizabeth Hinde, Danny M. Hatters

## Abstract

Mutations that cause Huntington’s Disease involve a polyglutamine (polyQ) sequence expansion beyond 35 repeats in exon 1 of Huntingtin. Intracellular inclusion bodies of mutant Huntingtin protein are a key feature of Huntington’s disease brain pathology. We previously showed that in cell culture the formation of inclusions involved the assembly of disordered structures of mHtt exon 1 fragments (Httex1) and they were enriched with translational machinery when first formed. We hypothesized that nascent mutant Httex1 chains co-aggregate during translation by phase separation into liquid-like disordered aggregates and then convert to more rigid, amyloid structures. Here we further examined the mechanisms of inclusion assembly in a human epithelial kidney (AD293) cell culture model and examined whether ribosome quality control machinery previously implicated in stalled ribosomes were involved. We found mHttex1 did not appear to stall translation of its own nascent chain and there was no recruitment of RNA into inclusions. However, proteins involved in translation or ribosome quality control were co-recruited into the inclusions (Ltn1 and Rack1) compared to a protein not anticipated to be involved (NACAD). Furthermore, we observed co-aggregation with other proteins previously identified in inclusions, including Upf-1 and chaperone-like proteins Sgta and Hspb1, which also suppressed aggregation at high co-expression levels. The newly formed inclusions contained immobile mHttex1 molecules which points to the disordered aggregates being mechanically rigid prior to amyloid formation.

## INTRODUCTION

Huntington Disease (HD) is an incurable and fatal neurodegenerative condition caused by dominant trinucleotide expansion mutations in exon 1 of the *Huntingtin* gene [1]. These mutations expand a polyglutamine (polyQ) sequence in the Huntingtin (Htt) protein to beyond a disease threshold of 35Q, which makes the protein become aggregation prone [2]. N-terminal mutant Htt fragments accumulate in intracellular inclusion bodies (inclusions) during disease progression, which represent a major hallmark of disease pathology [2-4].

The transgenic expression of the Htt exon 1 fragment (Httex1) in polyQ-expanded form is sufficient to produce a HD-like pathology in rodent and primate models, which is suggestive of these fragments mediating proteotoxicity [5-7]. The mechanism of toxicity remains to be unequivocally determined but it is thought to involve two distinct components; soluble and inclusion states of Httex1 [8]. Soluble states, which may include monomeric or small nanometer-sized oligomers of mutant Httex1 cause oxidative and mitochondrial stress and increase the risk of apoptosis in cell culture models of disease [9-12]. We previously suggested that the toxicity of the soluble forms of mutant Httex1 may involve a quality control feedback mechanism during translation involving stalled Httex1 nascent chains, which when unresolved triggers apoptosis [8]. Once inclusions form survival times are improved in cell culture models of disease, leading to a hypothesis that inclusion formation alleviates toxicity by sequestering the soluble toxic forms away from harm (reviewed in [13]). However, rather than returning the cell to a normal state of homeostasis, cells in culture with inclusions are metabolically quiescent and die at a delayed rate by a non-apoptotic necrotic mechanism [8]. This finding suggests a second level of toxicity from the inclusions distinct to that from the soluble states.

Here we sought to further investigate the molecular processes governing inclusion assembly in the context of the hypothesis that newly synthesized mutant Httex1 stalls at the ribosome, attracts ribosome quality control machinery to resolve the stress and when unresolved triggers the nucleation of initially disordered aggregates into liquid-like droplets that then over time convert to amyloid. Since our initial prediction that mutant Httex1 aggregates may arise through phase separation into liquid-like structures, two studies have since reported evidence supporting this mechanisms of action [14, 15]. Our findings suggest that nascent Httex1 does not stall on ribosomes during translation but there is an enrichment of machinery involved in ribosome associated quality control into the inclusions. In our hands and model, however, we found the early formed inclusions comprised immobile mutant Httex1 molecules with no evidence of liquid-like properties.

## METHODS

### DNA vectors and constructs

Human Httex1 and TC9-tagged Httex1 as fusions to fluorescent proteins were expressed in pT-Rex vectors with CMV-promoters as described previously [8]. The pFN21A-HaloTag constructs were purchased from Promega. The P2A stall construct was prepared as described previously including the Httex1 constructs [16]. The tandem P2A T2A constructs were made using T2A sequences from [17]. In essence the T2A sequence 5’ GGCGAGGGCAGGGGAAGTCTTCTAACATGCGGGGACGTGGAGGAAAATCCCGGCCCA was inserted after the P2A sequence of the existing Httex1(25Q) sequence in the pTriEx4 vector (GeneArt). The sequence of the derived vector is shown in **Table 1**. From this, the Httex1(97Q) and 20K variants were made by excising the gene fragments from the original stall reporter via NotI and BamHI restriction sites. The control linker was made by PCR amplication using forward (5’ GCGGCCGCTATGCCTGGACCTACACCTAGCG) and reverse (5’ GGATCCGCCGGTTTTCAGGCCAGGGC) primers and ligation into the P2A T2A Htt25Q Stall Reporter via the NotI and BamHI restriction sites.

**Table 1.**
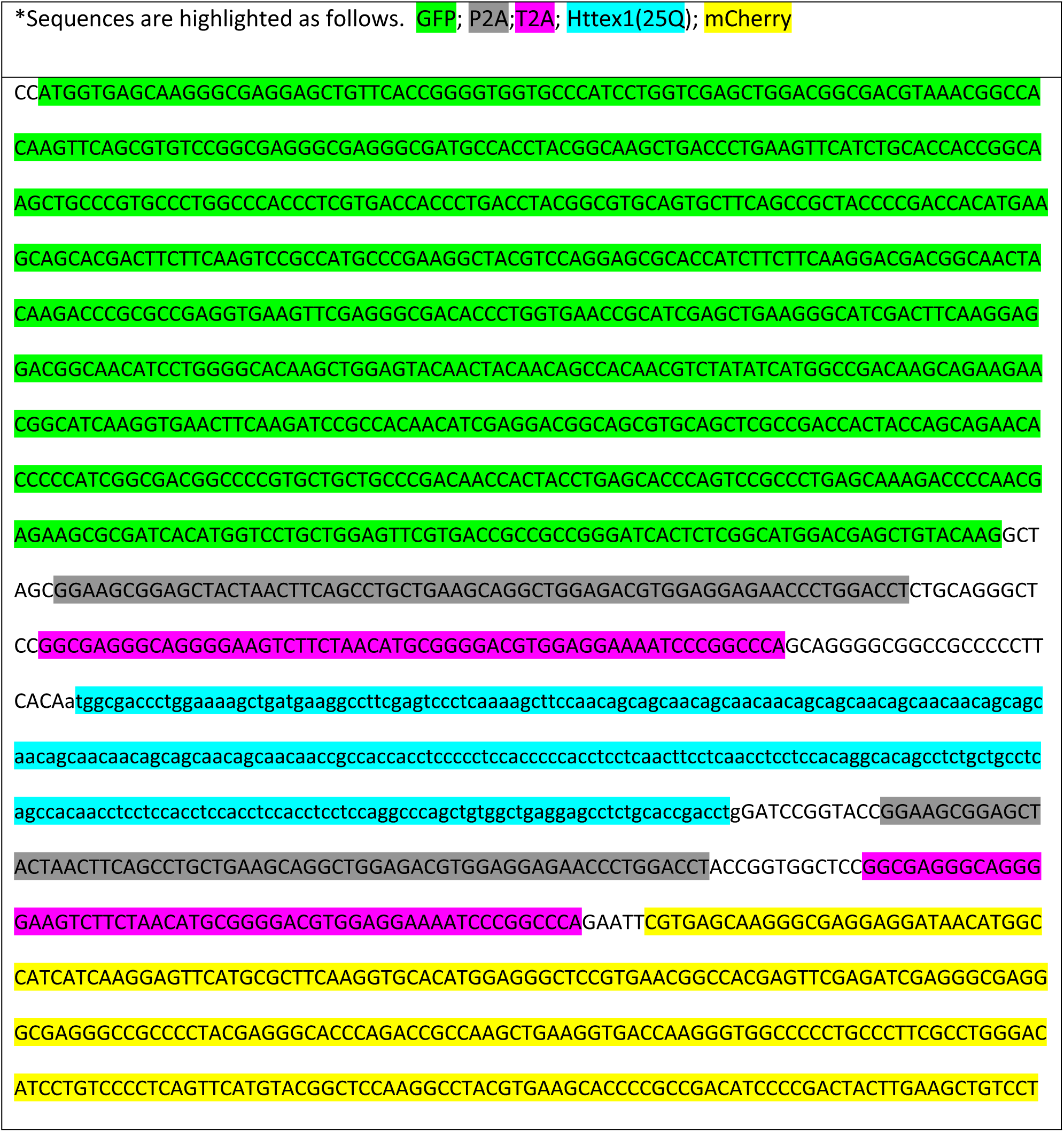
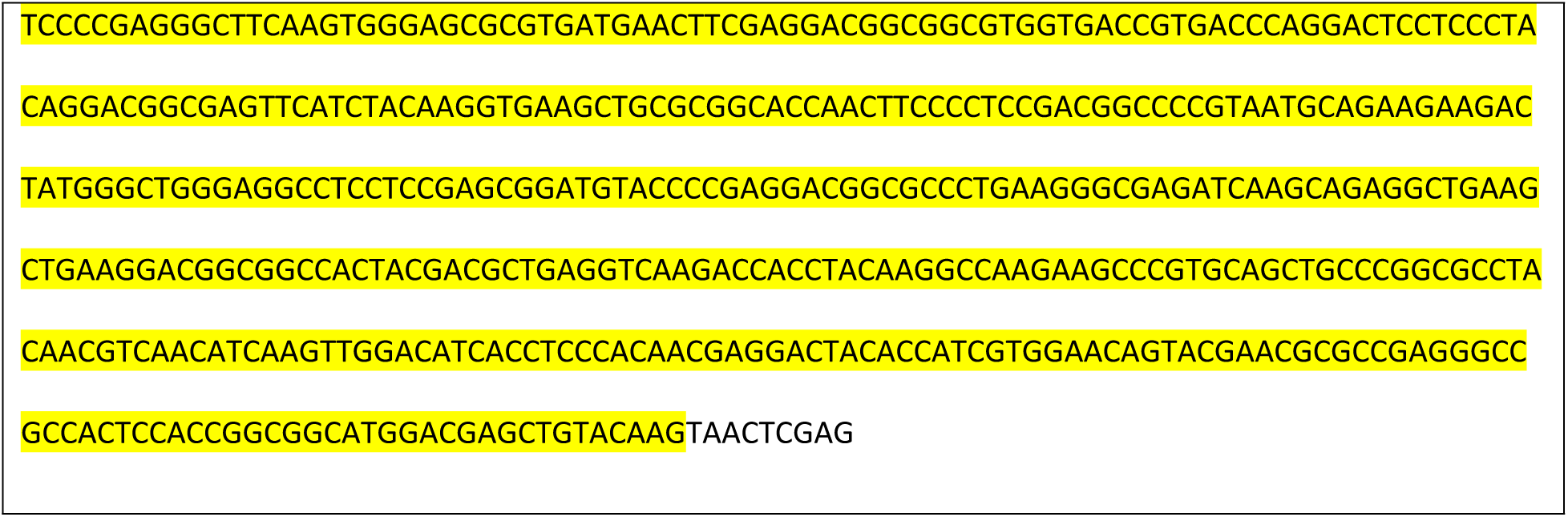
Sequence of the base stall construct *.

### Cell culture

All experiments were performed with the HEK293 cell line derivative AD293, which was maintained in DMEM supplemented with 10 % (w/v) fetal calf serum (FCS) and 1 mM glutamine in a 37°C humidified incubator with 5% v/v atmospheric CO_2_. For microscopy experiments cells were plated at 3 × 10^4^ cells per well in an 8-well µ-slide (Ibidi). For flow cytometry experiments cells were plated at 0.5 × 10^5^ cells in a 24-well plate. Cells were transiently transfected with the vectors using Lipofectamine 3000 reagent as per manufacturer’s instructions (Life Technologies), and media was changed 6 hours post transfection. For the HaloTag experiments, the transfection was done in a way so as to decouple the correlated expression of the two plasmids. Specifically, this was achieved by mixing the plasmids separately with Lipofectamine, before combining the lipofectamine:DNA complexes together to add to the cells.

### Western Blot

AD293 cells were transfected with P2A or P2A T2A stall constructs and harvested 24 hours post transfection. Cells were pelleted (200 *g*, 6 mins) and resuspended in RIPA lysis buffer (150mM NaCl, 50 mM Tris pH 8.0, 1% v/v IGEPAL, 0.5% v/v sodium deoxycholate, 0.1% v/v SDS, 250 U Benzonase and supplemented with cOmplete, EDTA-free Protease Inhibitor Cocktail pills (Roche)) and incubated on ice for 30 minutes. Lysate was matched for total protein by BCA kit (Thermofisher scientific, cat# 23225). 10 µg of total protein lysate was loaded on to an TGX Stain Free FastCast Acrylamide gel (BioRad, cat# 1610185) and transferred using an iBlot2 gel transfer device (Thermofisher scientific, cat# IB21001) and a PVDF iBlot2 transfer stack (Thermofisher scientific, cat# IB24001). The membrane was blocked with 5% w/v skim milk powder in phosphate buffered saline (PBS) for 1 hour at room temperature. Anti-GFP (Invitrogen, cat#A6455) and anti-Cherry (Abcam, cat#167453) were diluted to 1:10,000 and 1:2500 respectively in PBS containing 0.1% v/v Tween 20 and incubated for 1 hour at room tempterature. The secondary antibody, goat anti-rabbit HRP antibody (Invitrogen, cat#656120), was diluted 1:10,000 in PBS containing 0.1% v/v Tween 20 and incubated for 1 hour at room tempterature. HRP was detected by enhanced chemilumiscence.

### FlAsH staining

Cells were stained with FlAsH as described previously to demarcate HBRi from PBRi cells [8]. In essence, the ratio of Cerulean:FlAsH or mCherry:FlAsH fluorescence was determined and all cells with a ratio greater than one standard error from the mean were classified as HBRi whereas all cells with a ratio smaller than one standard error from the mean were classified as PBRi.

### RNA staining

Cells were stained for RNA using the Click-iT Plus Alexa Fluor647 Picolyl Azide Toolkit (Life technologies, cat#C10643) according to manufacture’s instructions. In short, 6 hours post-transfection 5-ethynyl uridine (Click-Chemistry tools, cat# 1261-10) was added to cells to a final concentration of 0.4 mM. 24 hours post-transfection cells were stained with FlAsH as described above. Following staining, cells were fixed with 4% w/v paraformaldehyde and permeabilised with 0.5% w/v Triton X-100 in PBS. Cells were then washed with 3% w/v bovine serum albumin in PBS followed by Click-It staining according to manufacturer’s instructions using a 1:4 ratio of CuSO_4_: Copper protectant. Nuclei were counter stained with Hoechst (1:1000) and imaged by confocal microscopy.

### HaloTag staining

After 6 hours post-transfection with the Halo-tagged constructs, TMRDirect ligand (Promega) was diluted 1:1000 in complete media and added to cells. For imaging, cells were fixed at 24 hours post-transfection with 4% w/v paraformaldehyde. Imaging positions were marked using the “mark and find” feature and then images captured on a Lecia TCS SP5. Cells were then stained with FlAsH as described above and the positions were re-imaged.

### Flow Cytometry

Cells were analyzed at high flow rate in an LSRFortessa flow cytometer, equipped with 488- and 561-nm lasers (BD Biosciences). 50,000–100,000 events were collected, using a forward scatter threshold of 5,000. Data were collected in pulse height, area, and width parameters for each channel. For Cerulean fluorescence, data were collected with the 405-nm laser and BV421 filter (450/50 nm). TMR fluorescence was collected with the 561-nm laser and using the PE-Texas Red filter (610/20 nm). All flow cytometry data were preprocessed with FlowJo (Tree Star Inc.) to assign live and inclusion-containing cells. Cells with inclusions were assessed by Pulse Shape Analysis [18]. The fluorescence intensity for individual cells in each channel was exported and analysed using custom python scripts (available at https://doi.org/10.5281/zenodo.3789864). Briefly, cerulean fluorescence was assigned to 20 logarithmic bins spanning the range of recorded intensities. Four logarithmic bins for HaloTag fluorescence were then assigned (none, low, medium, high) independently for each construct according to the maximum intensity for that construct. Finally, the proportion of cells containing inclusions within each combined cerulean-HaloTag bin was calculated.

### Immunofluorescence

Cells were fixed at 24 hours post-transfection with 4% w/v paraformaldehyde for 15 min at room temperature. Cells were then permeabilized with 0.2% v/v Triton X-100 in PBS for 20 mins at room temperature. Samples were blocked in 5% w/v bovine serum albumin in PBS for 1 hour at room temperature. Cells were stained with anti-GFP (1:300 dilution) (Invitrogen cat# A6455) diluted in PBS containing 1 % w/v bovine serum albumin and 0.05% v/v Tween 20 for 1 hr at room temperature. Samples were then incubated in goat anti-rabbit Alexa Fluro 647 (1:500) (Life technologies cat#A21244) diluted in PBS containing 1 % w/v bovine serum albumin and 0.05% v/v Tween 20 for 30 mins at room temperature.

### Fluorescence recovery after photobleaching

Cells were imaged at 37°C and 5% atmospheric CO_2_ on an Olympus FV3000 confocal laser scanning microscope through a 60X 1.2NA water immersion objective. mCherry fluorescence was excited with a 561 nm solid state laser diode. The resulting fluorescent emission was directed through a 405/488/561 dichroic mirror to remove laser light and detected by an internal GaAsP photomultiplier set to collect between 550–650 nm. 24 hours post transfection cells were FlAsH stained as described above and then imaged. For FRAP, a pre-bleached image was taken before half the inclusion was bleached for 5 seconds (10% laser power, 200 μs pixel dwell time). Recovery was then monitored by imaging the inclusions every minute for 21 minutes.

### Cell imaging and analysis

Images of live or fixed cells were acquired with a Leica TCS SP5 confocal microscope. Anti-GFP images were taken on the Zeiss LSM800 Airyscan. Images were extracted from proprietary formats using FIJI (v 2.0.0-rc-69/1.52p) equipped with the Bioformats plugin (v6.3.1). The resultant tiff images were then processed using custom scripts written for FIJI and python (source code available at https://doi.org/10.5281/zenodo.3789864). Briefly, in the case of RNA- and HaloTag-stained images, regions of interest (ROI) were manually assigned and the mean fluorescence intensity within each region exported. In the case of fluorescence recovery after photobleaching, bleached and whole-cell ROI were assigned via automatic thresholding of the FlAsH and mCherry channels respectively, while a circular background ROI with a nominal fixed radius (25 pixels in the images) was manually assigned for each image. Pixel fluorescence intensities for each ROI were exported for all timepoints post-recovery, and the bleached ROI determined by subtracting the unbleached pixel coordinates from the whole-cell ROI. The relative recovery was then calculated by dividing the background-corrected mean intensity in the bleached and non-bleached ROIs. Finally, in the case of anti-GFP antibody penetration, inclusions were identified by automatic thresholding of the cerulean intensity and the resultant ROI was scaled by 110% to yield the external inclusion boundary. The internal boundary was determined via automatic thresholding of the inverted anti-GFP fluorescence. The Euclidean distance between the centroid of the outer ROI and either the internal or external boundary pixels was then calculated, and the penetration measured as the difference in the mean distance of the internal and external boundaries.

### Statistics

The statistical tests are described in the figure legends and *P* values shown directly on the figures or coded as *, *P* < 0.05, **; *P* < 0.01, ***; *P* < 0.001; ****, *P* < 0.0001. Statistical tests were performed in Graphpad Prism v6.

## RESULTS

We previously postulated the presence of a translation-related quality control mechanism that clears aggregating or misfolding proteins emerging from the ribosome in cells lacking inclusions. Prior data has shown that polyglutamine (polyQ)-expanded Httex1 is more efficiently degraded than the wild-type counterpart, which is consistent with an elevated clearance mechanism at this step [19]. If polyQ-expanded Httex1 is engaging with quality control during synthesis we might anticipate this to lead to an interruption of translation rates. Furthermore we previously found, by proteomics, that Upf1 (Rent1), which plays a central role in non-sense mediated decay [20] was enriched in the inclusions [8]. This finding raises the possibility that the stalled constructs, should they arise, proceed to nucleate the aggregation process.

First to test for stalling, we implemented a translational stall assay in AD293 cells which are sensitive to the proteotoxicity of polyQ-expanded Httex1 [8]. The assay involves a reporter cassette containing two fluorescent reporters on each side of the peptide sequence to be tested for stalling (GFP at the N-terminus and mCherry at the C-terminus) [21] (**Fig 1A**). Each construct is encoded in frame without stop codons however the test sequence is flanked by viral P2A sequences, which causes the ribosome to skip the formation of a peptide bond but otherwise continue translation elongation uninterrupted [22]. Complete translation of the cassette from one ribosome will generate three independent proteins (GFP, test protein, and mCherry). However, should the ribosome stall during synthesis (such as through the previously established poly-lysine (20K) sequence used here as a control [21]), mCherry is produced at lower stoichiometries than the GFP. In our hands, we noted that there was a small fraction of protein synthesized reading through the P2A sequences (**Fig 1B**). Of particular note was the appearance of SDS-insoluble material in the stacking gel for the mutant polyQ-length form of Httex1 (97Q), which is indicative of SDS-insoluble mHtt products that arises from aggregation [3] (**Fig 1B**). Because fluorescence resonance energy transfer (FRET) from GFP to mCherry is anticipated to inflate the mCherry/GFP fluorescence ratio, and be particularly high in the aggregates, we sought to remove this confounding factor by redesigning the stall construct to have a more efficient skip sequence [17]. This involved tandem P2A and T2A sequences, which appeared to reduce the read through effect below detection by Western Blot (**Fig 1C**). The data shows Httex1 in wild-type (25Q) or mutant (97Q) form to have an insignificant difference in mCherry/GFP ratios compared to the control construct, although there was a very small significant difference between the 25Q and 97Q constructs. Collectively these data suggest that long polyQ sequences do not lead to ribosome stalling, or very small amounts, and therefore point to the proteotoxicity of soluble Httex1 most likely arising from other factors.

**Figure 1.**
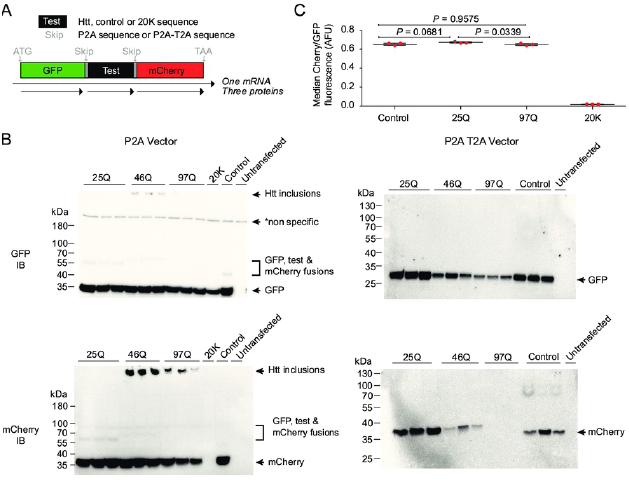
Translation of Httex1 is not stalled during synthesis at short or long polyQ lengths in AD293 cells. **A.** Schematic of the design of the stall reporter assays. GFP and mCherry sequences flank a test sequence and are expressed in frame. The 20K sequence is a positive stall control containing repeat AAA sequences as the test sequence. The Control linker is a non-stalling sequence for the test sequence. In the original assay, the P2A sequence causes the ribosome to skip the formation of a peptide bond leading to independent protein expression of the three components. If the ribosome stalls during translation of the test sequence, then the yield of mCherry decreases relative to GFP. In the modified assay, an additional alternative skip sequence (T2A) was added in tandem to the P2A to reduce the rate of readthrough **B.** Western blots of cells expressing the stall reporters probed with GFP or mCherry antibodies. Marker molecular weight masses are shown on the left. Products labeled on the right. **C.** mCherry:GFP fluorescence ratios measured by flow cytometry.

Next, we assessed whether the inclusions were enriched with RNA on the possibility that translational machinery, mRNA and ribosomes included, are coaggregated into the newly formed inclusions. For this we used a biosensor form of Httex1 (TC9-Httex1) we previously developed that enables us to demarcate early-formed inclusions from older inclusions [8, 18]. This biosensor involves a tetracysteine tag embedded near the start of the polyglutamine sequence that can bind to the biarsenical dye FlAsH when Httex1 is monomeric or forms a disordered aggregate in the early-formed inclusions. In contrast, when Httex1 is in an amyloid conformation and in the more mature inclusions, it is unable to bind [8]. Using a flow cytometry method called Pulse Shape analysis, cells containing Httex1 diffusely distributed (ni) or within inclusions (i) can be separated and then further divided into those Highly Biarsenical-Reactive (HBR) or Poorly Biarsenical-Reactive (PBR) [18, 23]. The result is the detection of cells with recently formed inclusions which are reactive to FlAsH (HBRi), and cells with mature inclusions which are PBRi [8]. HBRi and PBRi status can also be determined by microscopy [8].

To test for incorporation of RNA we used a Click-It RNA imaging approach, which involves adding 5-ethynyl uridine (EU) nucleoside to the media that then gets incorporated into newly synthesized RNA molecules and can be labelled with a fluorophore by click chemistry [24]. EU was rapidly incorporated into nuclear RNA pools and to a lesser extent the cytoplasm (**Fig 2A–B**). The levels of EU inside the inclusions was lower than the surrounding cytosol but was higher than background levels of fluorescence determined by control cells labelled with Alexa 647 in the absence of EU (**Fig 2C**). This suggests that at best there was a small incorporation of mRNA into the inclusions, however there was no difference between early-formed and later-formed inclusions in the amount of RNA. Hence, if nascent chain Httex1-ribosome complexes are major drivers of the first stages of aggregation, it is likely that this occurs after the 40S and 60S ribosome components have split, which would liberate the mRNA (and 40S subunit) from the stalled 60S-nascent chain complex [25, 26].

**Figure 2.**
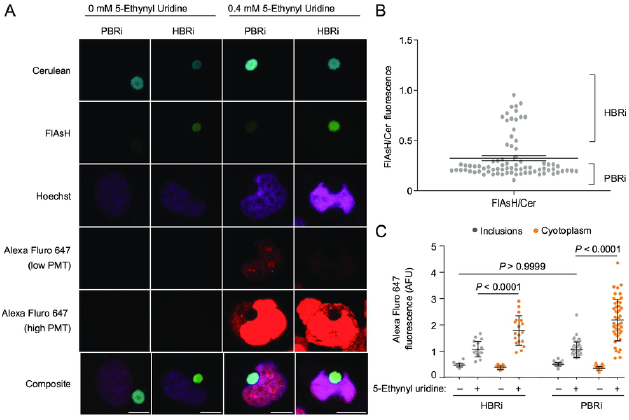
mRNA is not enriched inside Httex1 inclusions. **A**. Confocal images of AD293 cells transfected with TC9-Httex1(97Q)-Cerulean and stained with FlAsH 16 h after growing in nucleoside analogue 5-ethynyl uridine. Cells were stained with Hoecsht 33342 (nuclei stain), and Alexafluor 647 (RNA). Scale bar: 10 µm. **B.** FlAsH:Cerulean fluorescence ratios with ranges shown that was used to classify the HBRi and PBRi cells. Bar shows mean and SEM of individual inclusions. **C.** Quantitation of individual images of cells for intensity of Alexafluor 647 in one focal plane (measured by confocal microscopy). Data points indicate individual cells. Bars show means and SD. Differences were assesed by ordinary 2 way ANOVA.

We then examined whether components of the 60S ribosome and other candidate machinery involved in ribosome quality control (RQC) are enriched in the inclusions. The RQC system is responsible for monitoring partially synthesised proteins and labelling nascent chains that stall before reaching the stop codon for destruction [25, 26]. The RQC forms a stable complex with the 60S ribosome which triggers degradation of the nascent chain via ubiquitinylation [26]. The sequence of events include (1) splitting of the stalled ribosome; (2) assembly of the RQC and ubiquitinylation of the nascent chain; (3) extraction of the nascent chain and then degradation [25].

For these experiments, we co-expressed Halo-tagged candidate proteins previously suggested to be involved in ribosome quality control with TC9-Httex1(97Q), allowed inclusions to form and then stained for location of the candidate proteins via the HaloTags. This included Rack1 (Gnb2l1), which is a component of the 40S ribosome and is involved in translational repression [27, 28]. Rack1 has previously been reported to bind to Httex1, which in mutant form was suggested to interfere with protein translation [29], and is involved in initiating RQC by promoting ubiquitinylation of 40S ribosome subunits to resolve poly(A)-induced ribosome stalling [30]. We examined listerin (Ltn1) which is an E3 ubiquitin ligase that is recruited to 60S ribosome subunits close to the nascent chain and ubiquitinylates nascent chains that become stalled during synthesis [26]. We also investigated another protein not observed as enriched in inclusions [8] or predicted to be involved in this biology as a negative control, the α domain-containing protein 1 (Nacad). Nacad did not appear to colocalize with the inclusions based on HaloTag staining, yet both Ltn1 and Rack1 were enriched in the outer layer of the inclusions suggesting a specific enrichment of the RQC machinery to the aggregates (**Fig 3A** and **B**). The over-expression of these proteins did not appear to influence the formation of inclusions of Httex1(97Q) (**Fig 3C**).

**Figure 3.**
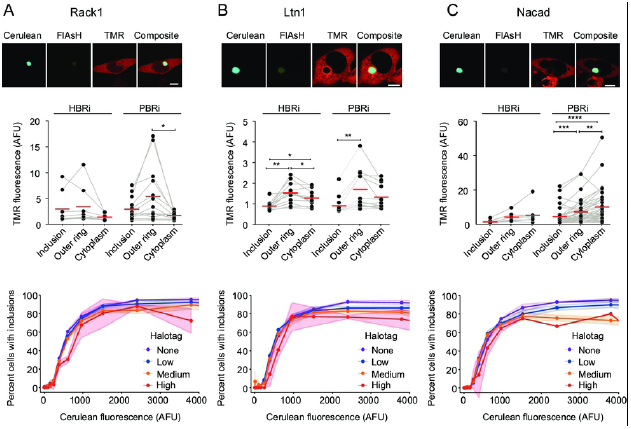
RQC proteins are recruited and enrich in the outer layer of inclusions whilst having no affect on inclusion formation. AD293 cells were co-transfected with TC9-Httex1(97Q)-Cerulean and Halo-tagged RQC proteins Rack1 (**A**), Ltn1 (**B**) and Nacad (**C**). HaloTag proteins were visualized by staining with TMRDirect ligand 24 hour after transfection (scale bar; 10 µm). Images (*upper panels*) were then quantified for enrichment of Halo-tagged proteins in inclusions (*middle panels*). The outer ring denotes the outer edge of the inclusion which was defined as the outer most area of cerulean fluorescence in the inclusions. Data points indicate individual cells, with lines connecting matched data. Means are shown as red dashes. Differences were assessed by repeated measures one way ANOVA with Tukey multiple comparisons test. Impact of HaloTag protein expression on Httex1(97Q)-Cerulean inclusion formation was determined by flow cytometry analysis (*lower panels*). Data indicates fraction of cells with inclusions, as measured by pulse shape analysis for cells categorized into different expression level bins (Htt levels, Cerulean fluorescence and Halo-tagged protein level, TMR fluorescence). The TMR was categorized into four evenly-spaced expression-level bins for each protein. Points and lines show means, shaded regions indicate SD.

To further probe the role of other proteins previously shown to be enriched in Httex1(97Q) inclusions [8] we overexpressed HaloTagged versions of proteins most enriched proteins in PBRi inclusions (Hspb1, Sgta, Upf1) and HBRi inclusions (Snu13, Rpl18). Hspb1 (Hsp27), which is a small heat shock protein involved in chaperone activity, appeared to enrich in the outer layer of Httex1(97Q) inclusions (**Fig 4A**). Furthermore, at high expression levels, Hspb1 lowered the potential of Httex1 to form inclusions which suggests it plays a role in mitigating (or reversing) aggregation (**Fig 4A**). Similarly, Sgta, which is a co-chaperone involved in the Bag6 system and ERAD also enriched to the inner and outer layer of Httex1 and had a more potent effect on suppressing aggregation of Httex1(97Q) at high co-expression levels (**Fig 4B**). Snu13, which is involved in pre-mRNA splicing, also was enriched in the inner and outer layers of the Httex1(97Q) inclusions and could suppress the aggregation of Httex1(97Q) (**Fig 4C**). Upf1 also enriched in the inner and outer edges of the inclusions but did not appear to affect the aggregation process of Httex1 (**Fig 4D**). Previously we found RPL18 as the most enriched protein in Httex1 inclusions by proteomics [8]. Halo-tagged Rpl18 was mostly present in the nucleus as punctate structures but small levels were seen in the cytosol (**Fig 4E**). Endogenous Rpl18 is anticipated to mostly reside in the nucleolus, ER and cytoplasm based on the Human Protein Atlas database [31]. We did not observe any evidence of enrichment of Halo-tagged Rpl18 with Httex1 inclusions (HBRi or PBRi). Furthermore, there was no evidence that the expression level of Halo-tagged Rpl18 affected the aggregation of Httex1 into inclusions (**Fig 4E**). However, the strong nuclear localization of the Halo-tagged Rpl18 suggests it might not be properly forming complexes with the ribosomes under these conditions.

**Figure 4.**
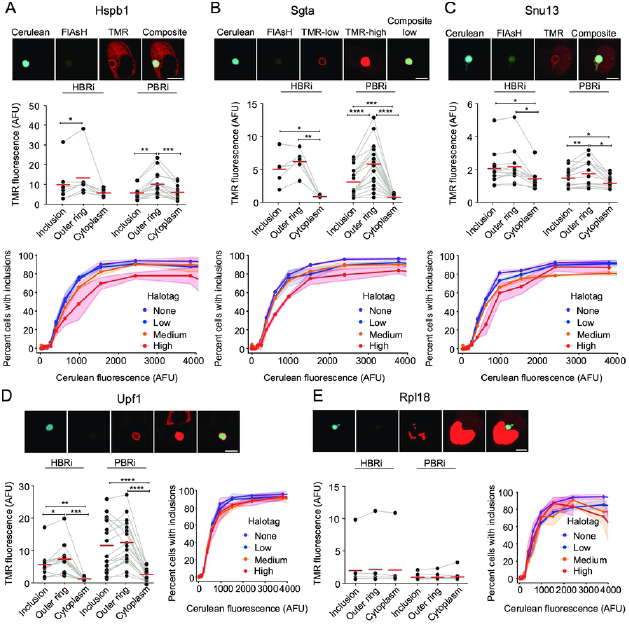
Recruitment patterns of proteins previously shown to be enriched in Httex1(97Q) inclusions and impact on inclusion assembly. AD293 cells were co-transfected with TC9-Httex1(97Q)-Cerulean and either chaperones Hspb1 (**A**) or Sgta (**B**), splicing proteins Snu13 (**C**) or Upf1 (**D**), or ribosome protein Rpl18 (**E**). Data is presented in the same manner as Fig 3. Scale bar; 10 µm. For proteins where cytosolic TMR staining could not been seen when other structures were not saturated a second ‘high PMT’ image was taken where the PMT was increased to a point when the cytoplasm could then be seen.

Our last set of experiments examined the diffusibility of mHttex1 in HBRi versus PBRi to determine whether the newly formed inclusions comprised of liquid-like mutant Httex1 aggregates as previously postulated [8, 14, 15]. For this experiment we performed fluorescence recovery after photobleaching on live AD293 cells expressing the TC9-Httex1(97Q)-mCherry and stained with FlAsH. After bleaching, both HBRi and PBRi inclusions did not recover over a period of 20 mins, which is consistent with a non-liquid aggregation state (**Fig 5A–B**). As a further probe for porosity we immunostained cells with a GFP antibody to test whether the antibody was able to penetrate the inclusions formed by TC9-Httex1(97Q) fused to GFP derivative Cerulean. The antibody formed a tight ring around the inclusions of both PBRi and HBRi, which suggested both inclusion states formed impenetrable aggregates (**Fig 5C**) and there was no statistical difference in penetration distance of the antibody staining between the HBRi and PBRi suggesting that inclusions form a dense, and immobile core structure quickly after formation (**Fig 5D**).

**Figure 5.**
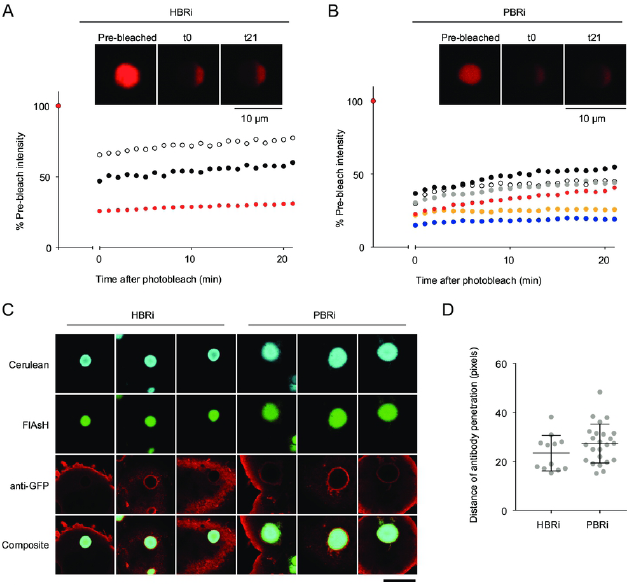
Httex1 molecules are immobile within early and later-formed inclusions. FRAP experiments tracking the recovery of mCherry fluorescence in regions of bleached HBRi (**A**) and PBRi (**B**) inclusions. Cells were expressing TC9-tagged Httex1 as a fusion to mCherry. Shown are representative images of inclusions before (pre-bleached), immediately after bleaching (t0) and at the endpoint (t21) following recovery. Plots show fluorescence recovery in the bleached region for individual inclusions. **C.** Inclusions in cells expressing Httex1 fusions to Cerulean are impervious to anti-GFP antibodies. Shown are representative images of anti-GFP stained inclusions for HBRi and PBRi (scale bar: 10 µm) **D.** Quantitation of antibody penetration distance into inclusions (pixels) from confocal images of immunostained inclusions in situ in cells. Individual inclusions are shown as well as means ± SD. There was no statistical difference as assessed by a two tailed t-test (*P*=0.1641).

## DISCUSSION

Our data here suggests that expression of Httex1, at wild-type or mutant polyQ lengths, does not stall on the ribosome during translation. As we wrote this manuscript a study was published that suggested Htt has a physiological function of binding to ribosomes and slowing the speed of ribosome translocation of many target mRNAs [32]. Intriguingly, mutant Htt further slowed protein synthesis rates [32]. These data suggest that Httex1 may be slowing translation in trans rather than in cis (through nascent Httex1-driven stalling). This data therefore raises the possibility that mutant Httex1 does not itself stall during synthesis but upon aggregation can sequester proteins similarly involved in regulating ribosome translocation, and potentially those involved in ribosome quality control, into mutant Httex1 inclusions. This mechanism likely would involve the co-aggregation of Httex1 with the endogenous full length Htt that exerts the physiological activity. It is important to note that our constructs contained mixed CAG and CAA codons of glutamine, which is different to the mostly homogenous CAG repeats seen in the human disease [1]. Other studies of polylysine encoded by repeating AAA codons stall much more effectively than polylysine encoded by mixed AAG and AAA lysine codons, or just AAG codons[33, 34]. The mechanism of stalling was explained in this case by contributions from both RNA and protein [34].

Previously it was shown that inclusions formed in yeast constitutively expressing mHtt(72Q)-GFP had a diffusible core suggestive of a liquid-like state [15]. Also it has been shown that Httex1 can form droplets in vitro and liquid-like states in cells [14]. We did not observe any evidence of liquid-like states in early or late-formed inclusions. It remains possible that the early-formed inclusions detected by our dyes have already solidified into a gel state but remain disordered by the time we assessed them. It also remains possible that the structures observed in other studies are distinct to what we observed.

Of the proteins that were co-aggregated Sgta, Snu13 and Upf1 were mildly enriched inside the inclusion whereas Rack1, Ltn1 and Hspb1 were only or more extensively enriched on the outside edge of the inclusion. Previously it was shown that Hspb1 can form molecular condensates, which raises the possibility of a mixed phase separation process with polyQ that may explain some of the co-aggregation mechanism [35, 36]. Nonetheless, the overexpression of the proteins involved in ribosome quality control did not alter the aggregation propensity. This result is more consistent with them playing non rate-limiting roles if they are involved in aggregation or clearance, or indeed acting as bystanders that do not play a critical role in mediating inclusion formation but are co-aggregated.

In conclusion, our data suggests that nascent chains of mutant Httex1 emergent from the ribosome are unlikely to stall and therefore unlikely to drive inclusion formation as stalled entities. However, given that we did see some ribosome-associated proteins co-aggregating as well as other proteins we previously identified as enriched in inclusions by proteomics, it remains possible that newly synthesized nascent Httex1 contributes to the aggregation process substoichiometrically by nucleating further association of post translated pools of Httex1. Alternatively, it is possible that aggregation of mHttex1 can co-aggregate with endogenous Htt that is engaging in a physiological function of regulating ribosome translocation rates, and thereby drawing translation machinery into the inclusions in trans. Both contexts are consistent with other reports of pre-existing pools of Httex1 monomer and small oligomers being quickly absconded into the inclusion once they form [37].

## Competing interests

No competing interests were disclosed.

## Grant information

This work was funded by grants and fellowships from the National Health and Medical Research Council Project to D.M.H. (APP1184166, APP1161803, APP1102059, APP1154352).

